# Bisbee: A proteomics validated analysis package for detecting differential splicing, identifying splice outliers, and predicting splice event protein effects

**DOI:** 10.1101/2020.08.13.250167

**Authors:** Rebecca F. Halperin, Apurva Hegde, Jessica D. Lang, Elizabeth A. Raupach, C4RCD Research Group, Christophe Legendre, Winnie S. Liang, Patricia M. LoRusso, Aleksandar Sekulic, Jeffrey A. Sosman, Jeffrey M. Trent, Sampathkumar Rangasamy, Patrick Pirrotte, Nicholas J. Schork

## Abstract

Here we present a novel statistical approach to splicing outlier and differential splicing detection, implemented in a software package called Bisbee. We leverage Bisbee’s prediction of protein level effects to benchmark using matched RNAseq and mass spectrometry data from normal tissues. Bisbee exhibits improved sensitivity and specificity over existing approaches. We applied Bisbee to confirm a pathogenic splicing event in a rare disease and to identify tumor-specific splice isoforms associated with an oncogenic splice factor mutation. We also identified common tumor associated splice isoforms replicated in an independent dataset, demonstrating the utility of Bisbee in discovering disease relevant splice variants.

## Introduction

Alternative splicing has been shown to play an important role in normal cellular processes as well as a wide range of pathogenic processes underlying many different diseases [1,2]. For example, global dysregulation of splicing, as well as mutations in genes regulating splicing, such as SF3B1, have been observed in a variety of tumors [3,4]. In addition, the results of genome wide association studies (GWAS) focusing on common chronic conditions have identified a number of disease-associated variants that influence splicing, suggesting a role for alternative splicing in mediating many common diseases [5,6]. Furthermore, highly penetrant variants that affect splicing have been classified as pathogenic in a number of monogenic disorders [7]. The detection of disease relevant splice alterations (*i.e.,* splice variants) is not trivial, as there are hundreds of thousands of annotated splice sites in the human genome. In addition, there is also great potential for the emergence of novel unannotated sites at countless locations in the genome. This suggests a need for robust statistical methods for detecting and quantifying differential splice events in comparative studies in health and disease. We have developed a novel statistical framework for differential splicing and splice outlier detection. We have implemented these methods for RNAseq data splicing analysis in a package called Bisbee (Figure 1). Bisbee also provides protein-level splicing effect predictions. We validated these predictions and benchmarked our statistical methods using normal tissue samples with both RNAseq and mass spectrometry data [8].

**Figure 1.**
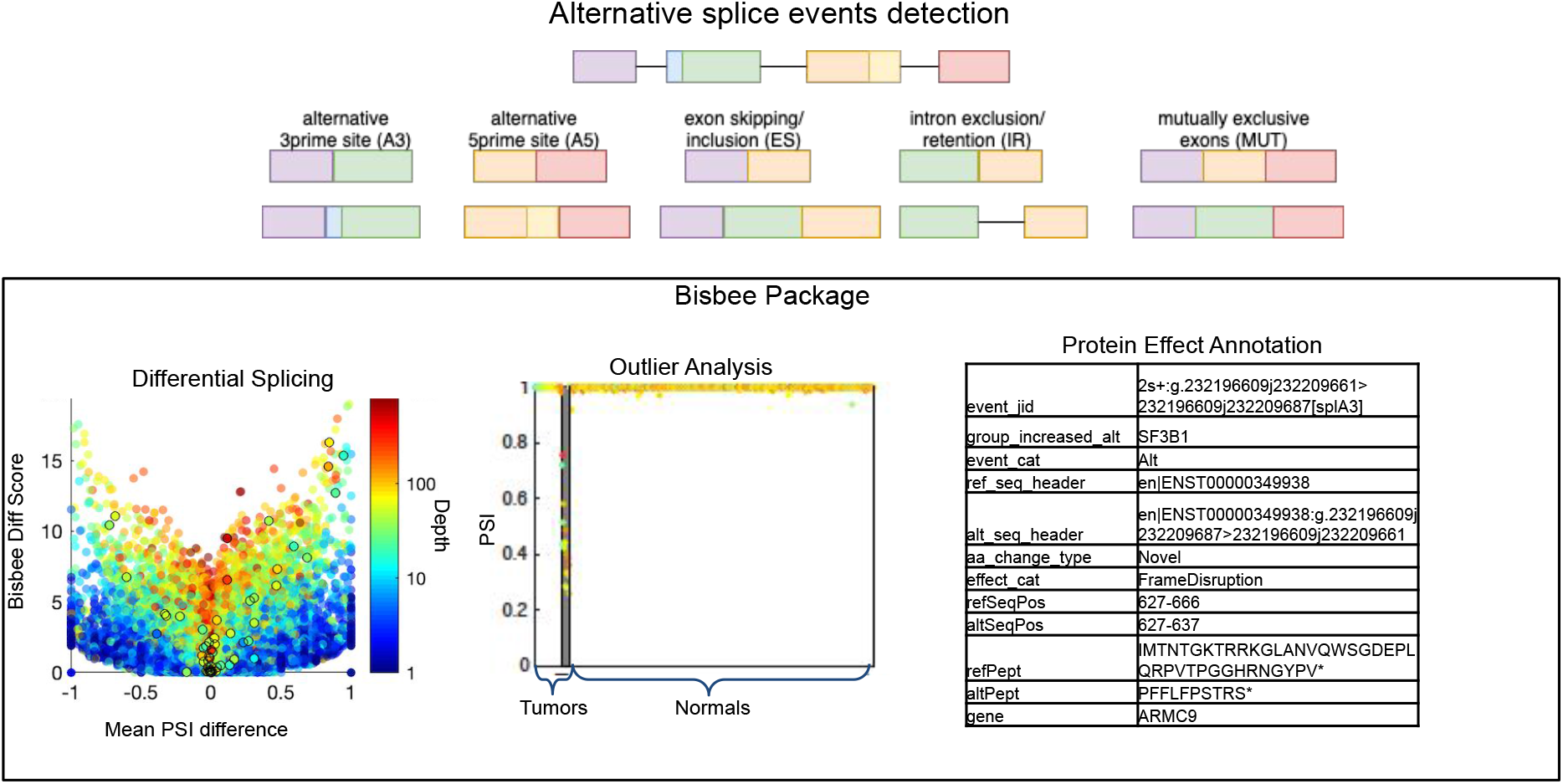
Method Overview. Five types of alternative splicing events are detected by SplAdder. For each event, two alternative splice isoforms are considered. Bisbee takes the read counts supporting each isoform in each sample and performs differential splicing or outlier analysis. As illustrated in the volcano plot on the left, Bisbee Diff is able to detect high coverage events with subtle differences in percent spliced in (PSI) as well as low coverage events with large differences in PSI. As illustrated in the center plot, the Bisbee outlier test also takes into account the differences in PSI and the coverage of the event. Each dot represents a sample, with tumors on the left and normal tissues on the right. The samples are sorted by outlier score within each set on the x-axis, the PSI is plotted on the y-axis, and the color represents the depth of coverage of the event in the sample. The dots within the grey stripe pass the outlier score threshold. Bisbee also annotates protein level effects and example output is shown on the right.

Alternative splicing analysis consists of three main steps: detection, statistical comparison, and effect prediction. Here we leverage an existing tool for detection and implement new methods for statistical analysis and effect prediction steps. Software packages for detecting splicing alterations may be broadly broken down into two categories: those that only identify events found in annotated transcripts such as ballgown [9], MISO [10], rMATs [11], and SUPPA2 [12], and those that additionally detect novel splice events, such as ASPLI [13], SplAdder [14], SGSeq [15], LeafCutter [16], and MAJIQ [17]. As aberrant splicing in disease states may result in novel transcripts, we sought to identify and extend the capability of a tool that can identify novel splice events. Novel splice events may reflect a well-annotated gene exhibiting previously unknown alternative splicing that occurs in diseased-tissue as opposed to normal tissue. We chose SplAdder for basic splice event detection because it has demonstrated utility in a large pan-cancer study and can enable comparisons to large sets of normal tissues from GTEx without requiring access to raw GTex data [14]. SplAdder also has advantages in terms of its modularity, facilitating analysis of large datasets in a cluster computing environment, and it also reports splice event coordinates in a straightforward manner.

Several splicing analysis packages include functions for testing differential splicing between two groups (e.g., diseased and non-diseased tissues), including ballgown [9], ASPLI [13], and SplAdder [14]. These typically use a generalized linear model and treat the overall expression level of the gene as a covariate to normalize expression differences that may confound the detection of splice isoform frequency differences. However, a more straightforward approach would be to explicitly test the difference in the ‘percent spliced in’ (PSI), and therefore obviate the need to normalize for library size or expression level. As PSI values are bounded between zero and one, they violate normality assumptions, *e.g.,* of a Student’s t-test, precluding its use to test the difference in PSI between sample groups. Similarly, the low number of biological replicates in most RNA-seq experiments limits the use of Wilcoxon’s test and affects the ability to measure the variance of a given event. To alleviate these challenges, we developed a novel approach to differential splicing analysis, which applies a beta binomial model in which the read counts supporting the splice event are modeled as a binomial distribution, and the underlying distribution of the PSI of a particular splice event in the samples of interest are modeled with a beta distribution. The binomial model captures the noise in the technical measurement of the PSI value due to the depth of coverage at the splice event without needing to rely on replicates, and the beta distribution models the biological variation in the splicing. Beta binomial models are used in many DNA sequencing variant calling strategies to model the distribution of reads supporting the existence of reference and alternate alleles [18–21]. Here we consider that each splice event has two alternate alleles, such as an exon included versus skipped in a particular gene, or the use of a first or second splice site for an exon (Figure 1). Our proposed approach is most similar to the strategy implemented in the ‘LeafCutter’ program, but LeafCutter does not use a binomial model, but rather a multinomial model, to test for differences in intron usage within a region [16]. Rather than explicitly reporting splice event types such as exon skipping or alternate splice site usage, LeafCutter reports clusters of alternative splicing. We chose to work with defined splice event types for improved interpretability and potential for insight into the mechanism of splice dysregulation.

In a precision medicine context, it is important to identify splice events that are verifiably occurring in an individual compared to a large reference set of samples rather than comparing two groups of samples for differences. This outlier analysis may be useful to identify disruption of splicing due to somatic mutations or expression of known tumor specific splice isoforms in an individual’s tumor [22,23]. Outlier analysis may also be used to identify splice variant-induced antigens in a target individual’s tumor that do not exist in normal tissues [3,23–25]. In addition, some rare Mendelian disorders are caused by variants that disrupt splicing. When searching for the causal variant in a rare disease, one is typically limited to studying a single family with a small number or even just one affected individual. Therefore identification of splice disrupting variants may require comparing an individual transcriptome to a set of reference samples. We are aware of only one tool designed for detecting splicing outliers in individual genomes, LeafCutterMD [26], which does not enable protein effect prediction.

Predicting the protein level impact of a splice variant is critical for understanding the biological implications and potential mechanisms underlying disease states, yet most alternative splicing analysis packages do not incorporate an effect prediction step or methodology. Splice variants may result in truncations or deletions at the protein level that result in a loss of protein function. Alternative splicing may also result in alternative protein isoforms that exhibit qualitative differences in function. For example, an alternate splice site in the BCL2L1 gene gives rise to different protein isoforms that have opposing effects on apoptosis [27]. Alternative splicing may also give rise to novel protein sequences in a cancer cell that could be recognized by the immune system [3,23–25]. In this light, genomics, transcriptomics, and proteomics are being used together more often in an effort to more fully characterize phenotypic effects resulting from genomic alterations and pathway dysregulation [28–30]. Predicting splice variant protein sequences in such studies would enable the creation of personalized predicted proteomes for mass spectra matching and enable more comprehensive and patient-specific proteomics analysis. Additionally, protein sequence prediction may provide insight into the functional effects of the splice alterations, as one could readily determine how much of the protein sequence is affected, which domain(s) are impacted, and make use of other in silico prediction tools for more detailed insight into splice variant effects on protein structure.

We have developed a suite of tools to identify and characterize splice events and variants called ‘‘Bisbee.’ Bisbee can identify differentially spliced variants between two conditions via the ‘Bisbee Diff’ test, can identify rare or individual-specific splice events via the ‘Bisbee outlier’ test, and can annotate splice events and predict protein isoform sequences via ‘Bisbee prot’ (see Methods). We evaluated the performance of these tools using various datasets. In order to compare different strategies for identifying and characterizing splice events and variants we developed a ‘truth set’ with splice events validated through the detection of corresponding protein isoforms. This truth set was generated using mass spectrometry and RNAseq data on a set of normal tissues from Wang et al. [8]. We identified several other splice variant analysis tools to consider for benchmarking against Bisbee. However, only a few of them provide utilities for predicting the effect of splice alterations at the protein level, which would be necessary for use with our mass spectrometry truth set. Using real data with complementary measurements for generating a truth set provides a more robust framework for benchmarking and validation than previous studies that have primarily relied on simulated data. In addition, simulating splice variant data requires making assumptions about the underlying distribution of splice events, the magnitude of the expected splicing differences in different biological conditions, and the nature of measurement error and other sources of noise. Our truth set takes advantage of the naturally occurring differences in splicing between different tissues [30]. We performed differential splicing analysis between several pairs of normal tissues and looked for enrichment of tissue-specific protein isoform expression to evaluate the results. Similarly, we used a set of gastrointestinal (GI) tissues as a reference set and looked for outliers in non-GI tissues, and evaluated evidence for protein level concordance among reported events.

## Results

### Predicted splice isoforms are detected at the protein level

In order to validate the existence of proteins/peptides corresponding to splice variants, we leveraged a dataset from Wang *et al.,* which includes paired RNA-seq and proteomics data from normal tissues [8]. In this validation dataset, SplAdder identified 268,791 total splice events, of which 125,683 were predicted by Bisbee to be protein coding. Of these protein coding events, we were able to detect at least one isoform-specific peptide for 74,817 splice events, including 2,033 generating novel sequences. Peptides supporting both protein isoforms were detected in 1,667 of the protein-generating events, including 1,144 generating novel sequences. Figure 2 illustrates the proportion of these events where both isoforms were detected with respect to the event type and event effect. All of the categories with at least 100 splice events have at least two of those events with confirmed protein level expression. The event categories that generate longer stretches of altered sequence have higher proportions of protein level detection as expected (Supplemental Figure 1). We observed 330 events showing tissue specific detection patterns at the protein level, and these were used for benchmarking and validation.

**Figure 2.**
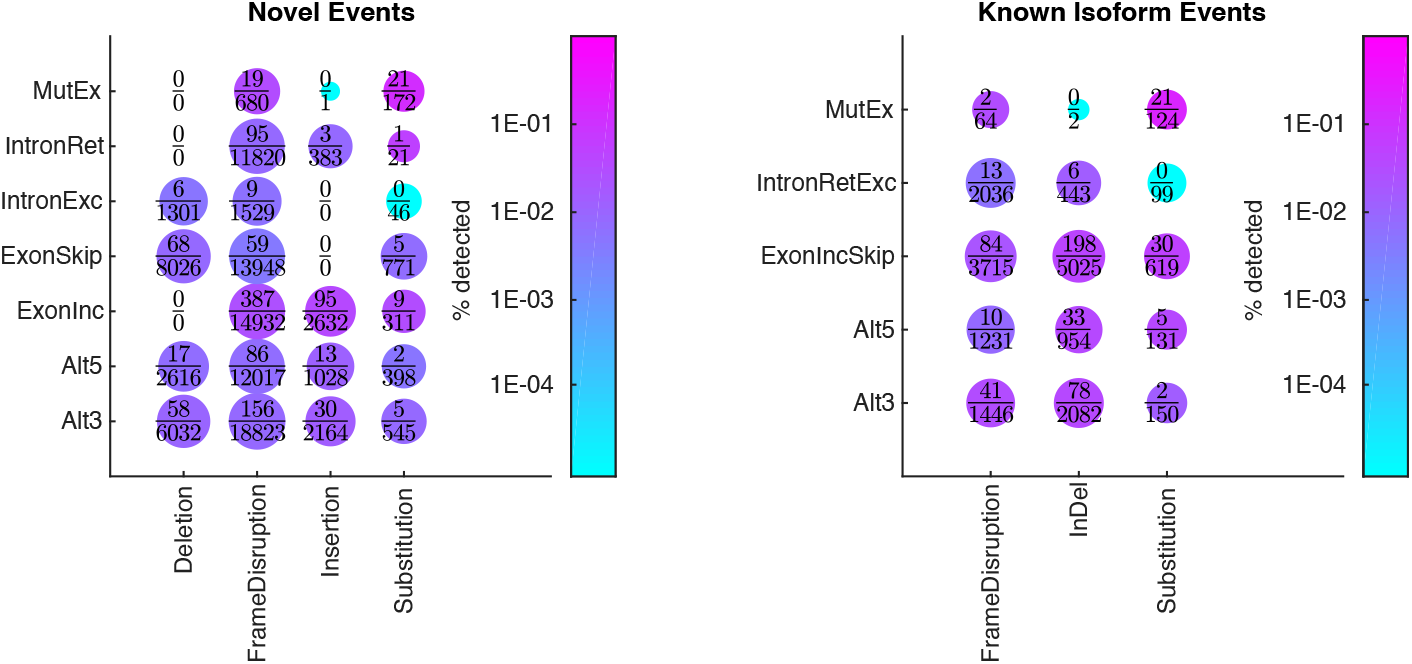
Bisbee detects many types of splice events with confirmed protein expression. The alternative splicing events generating protein coding sequences are categorized by event type and protein effect. Events where both isoforms exist in annotated transcripts are shown on the right and events where one isoform is not found in annotated transcripts are shown on the left. The size of each dot is proportional to the number of predicted sequences in each category from the normal tissue RNAseq dataset and the proportion of events confirmed by mass spectrometry data is indicated by color. ‘Alt’ indicates alternative 3 or 5 prime splice sites, ‘Exoninc’ indicates exon inclusion, ‘ExonSkip’ indicates exon skipping, ‘IntronExc’ indicates intron exclusion, ‘IntronRet’ indicates intron retention, and ‘MutEx’ indicates mutually exclusive exons. For novel events, exon skipping/inclusion, intron retention/exclusion, and insertions/deletions are categorized by the effect on the novel isoform. Frame disruption events include those that shift the reading frame as well as those which result in the use of a different stop codon such as in an intron or untranslated region (UTR).

### Bisbee Diff more accurately detects differentially spliced isoforms

The beta binomial differential splicing test implemented in Bisbee has one parameter that requires tuning. In order to avoid overfitting, we reserved the tissue-specific protein isoforms dataset from Wang et al. to compare the accuracy of differential splicing methods. GTEx was used for parameter optimization and threshold selection. We compared the distribution of the test statistic for the Bisbee Diff test between sets of samples from the same tissue versus different tissues using different values of the *ω_M_* parameter. The percentage of events passing a given threshold in the ‘different’ versus the ‘same’ comparison is used as an indicator of the specificity of the test, while the percentage of events in the different comparisons passing the thresholds is used as an indicator of sensitivity. Setting the *ω_M_* parameter to 200, and using a log likelihood ratio (LR) threshold of 8 provides optimal enrichment of splice events detected as different between different tissues compared to splice events detected as different between samples from the same tissue. (Supplemental Figure 2a).

We identified 281 instances of protein expression-confirmed isoform switches over six pairwise tissue comparisons, which represent 196 unique isoform switch events. For comparison, SplAdder’s test module was run as an example of the generalized linear model approach. As a simple approach, a t-test on the PSI values was pursued both using all of the PSI values regardless of depth and only including PSI values with a sequencing read depth at the position of greater than 10. To evaluate these methods, we compared the total number of events passing a given threshold to the number of protein confirmed events passing the threshold. The Bisbee Diff method consistently found higher enrichment of confirmed events out of total events passing a threshold (Figure 3a). Counts of the number of confirmed versus total events are also shown in Table 1. In order to see how the magnitude of PSI differences and the read depths of the events influence the performance of each of the differential splicing tests we made a volcano plot of the brain versus small intestine comparison (Figure 4).

**Figure 3.**
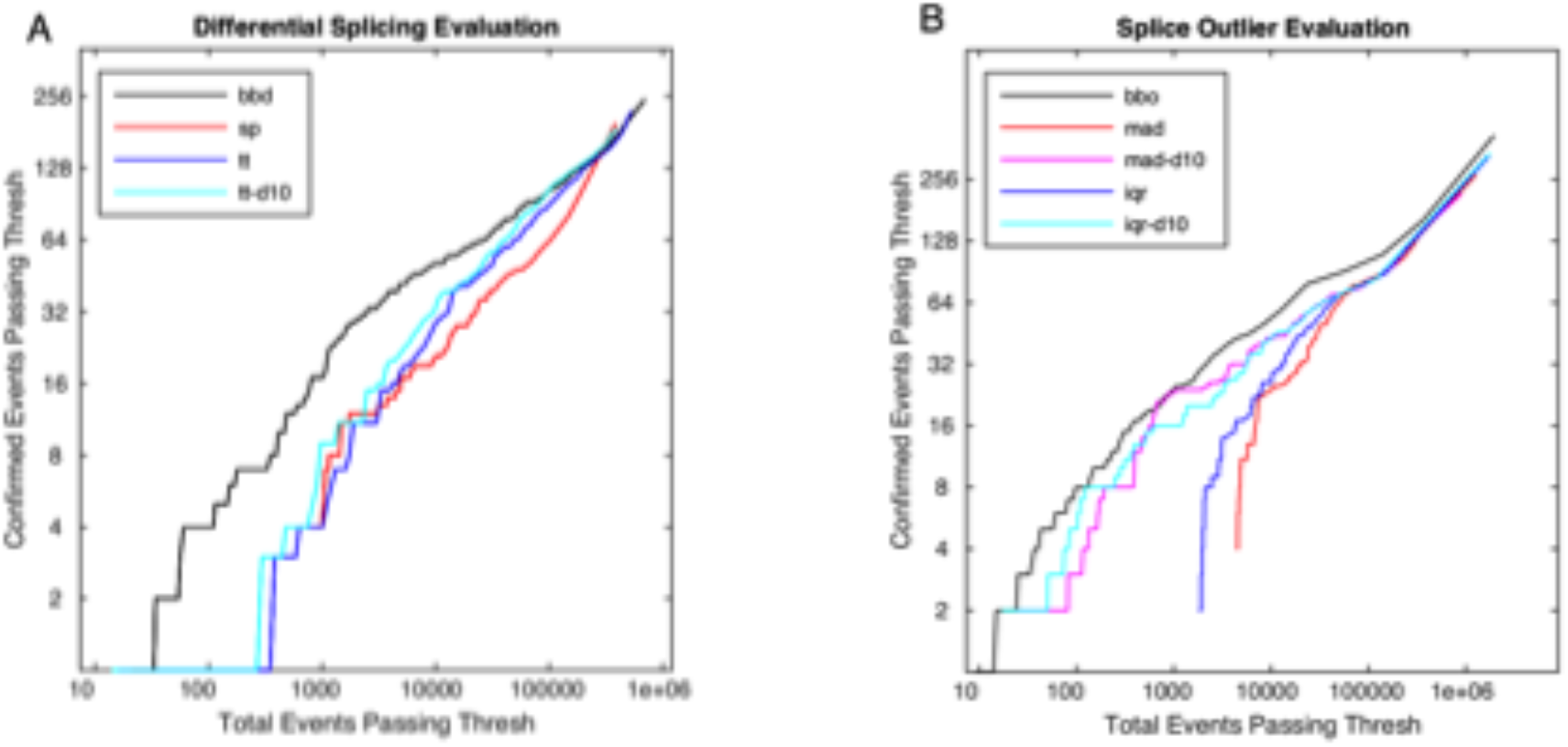
Bisbee detects protein expression confirmed splice events with high sensitivity. a) The number of events with protein expression evidence of differential splicing is plotted against the total number of events passing the threshold for four different differential splicing methods: beta binomial (bbd – black), SplAdder’s test (sp – red), t-test on all PSI (tt – blue), t-test on PSI with depth>10 (tt-d10 – cyan). b) The number of mass spectrometry confirmed outlier events is plotted against the total number of events passing the threshold for five different methods: bisbee outlier (black), median absolute deviation (red), median absolute deviation with depth>10 (magenta), interquartile range (blue), and interquartile range with depth>10 (cyan).

**Figure 4.**
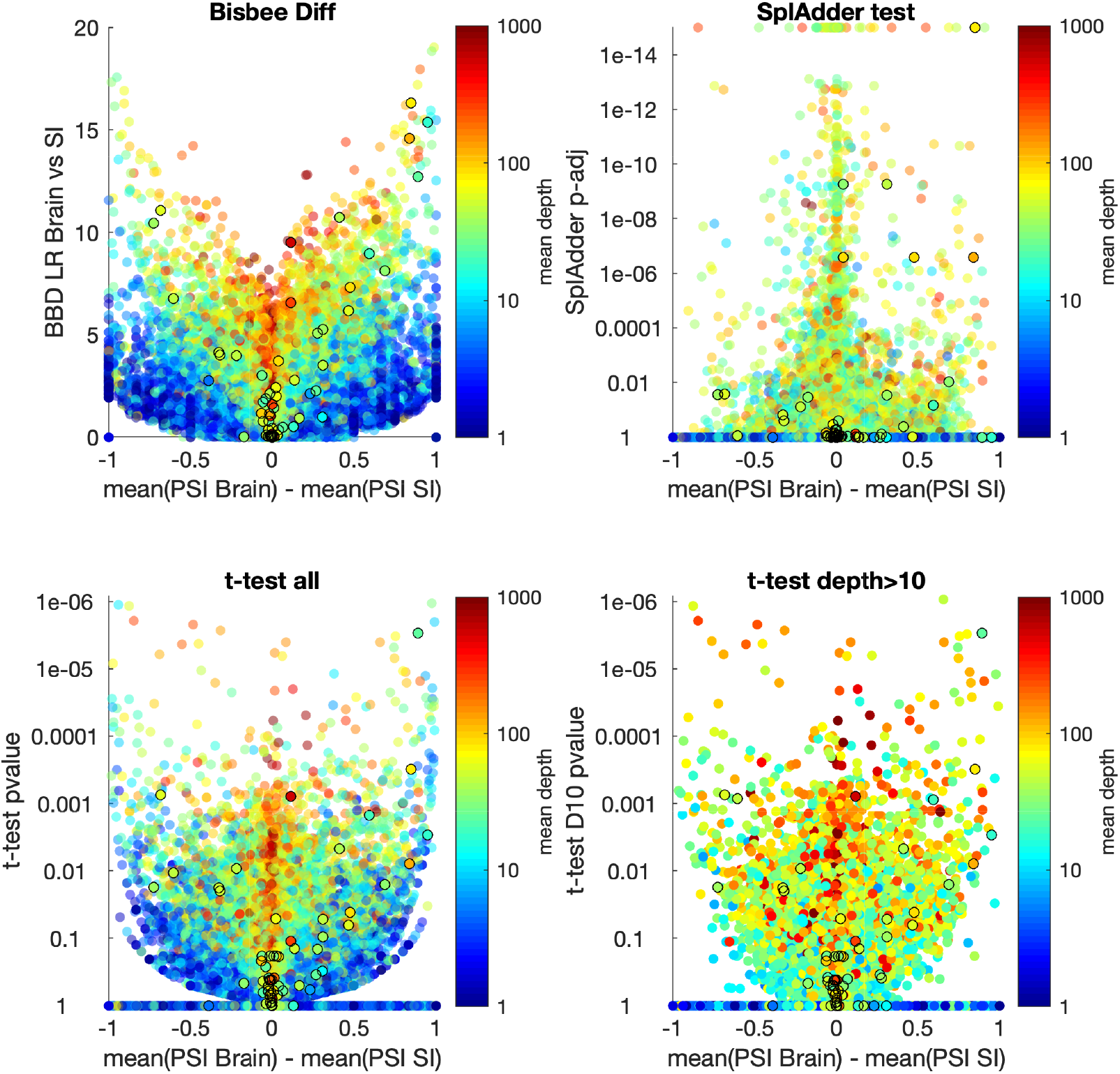
Volcano plots of brain versus small intestine for four differential splicing methods. Differential splicing results for the brain versus small intestine comparison from the Wang et al. dataset. The Bisbee Diff LLR (a), SplAdder test adjusted p-value on a log scale (b), or t-test p-value on a log scale (c and d) are plotted against the difference in mean PSI between the brain and small intestine samples. Points are colored by the mean read depth covering the event on a log scale as indicated by the color bar. Events with mass spectrometry confirmed tissue specific protein expression are indicated by black circles.

**Table 1.** Differential splicing benchmarking summary

| Name | 1st threshold | Confirmed pass 1st | Total pass 1st | 2nd threshold | Confirmed pass 2nd | Total pass 2nd |
| --- | --- | --- | --- | --- | --- | --- |
| Bisbee Diff | $lr > 8$ | 28 | 1760 | $lr > 12$ | 7 | 196 |
| SplAdder Test | $p < 0.01$ | 35 | 25848 | $p_{adj} < 0.05$ | 28 | 16717 |
| T-test all | $p < 0.01$ | 25 | 8645 | $fdr < 0.05$ | 1 | 127 |
| T-test D10 | $p < 0.01$ | 26 | 6448 | $fdr < 0.05$ | 1 | 103 |

**Table 2.** Outlier test benchmarking

| Name | 1st threshold | Confirmed pass 1st | Total pass 1st | 2nd threshold | Confirmed pass 2nd | Total pass 2nd |
| --- | --- | --- | --- | --- | --- | --- |
| Bisbee Out | 10 | 37 | 2947 | 14 | 28 | 1674 |
| MAD | 20 | 32 | 19371 | 40 | 25 | 10859 |
| MAD-D10 | 20 | 32 | 3883 | 25 | 26 | 2712 |
| IQR | 10 | 36 | 14146 | 20 | 26 | 8178 |
| IQR-D10 | 8 | 36 | 7694 | 10 | 34 | 5597 |

### Bisbee outlier more accurately detects splice outliers

The Bisbee outlier detection method parameter *β_M_* was optimized using GTEx data. The percentage of outlier scores passing a threshold for models trained on the same tissue was compared to the percentage passing for matching tissue models. We found a *β_M_* value of 80 provides the best enrichment of different tissue outliers with a log likelihood (LL) cutoff of 10 (Supplemental Figure 2b). We used these values for benchmarking on the Wang *et al.* dataset with matching proteomics data [8]. We used a set of GI tissues as the reference set and detected outliers in three other tissues. We identified 140 outlier events across the three tissues, which represents 134 unique outlier events. Since we are not aware of another tool that is able to detect splice outliers and generate predicted protein sequences, we implemented two simple methods using the distribution of PSI values in the reference dataset. The first simple outlier method uses the median absolute deviation (mad) and the second using the interquartile range (iqr) of the PSI values. For both of these methods we performed the analysis both with using PSI values for all data points as well as using only PSI values for data points with a depth greater than 10. The Bisbee outlier method detected more proteomics-confirmed events for similar numbers of total events passing the same score threshold (Figure 3b).

### Case study: detection of splice event in rare disease

In order to examine the utility of the Bisbee package for research and clinical applications, we analyzed disease-causing splice mutation in the nuclear-encoded mitochondrial methionyl-tRNA formyltransferase (MTFMT) [31–33]. We previously identified homozygous mutation (c. 626 C>T) in the MTFMT gene in three children from two unrelated families (Clinvar Accession#VCV000039827.4) with Leigh syndrome and combined oxidative phosphorylation (OXPHOS) deficiency. The MTFMT mutation c. 626 C>T in the coding region resulted in a Ser209Leu (S209L) amino acid substitution, which is likely a non-pathogenic event. However, c.626 C>T is predicted to generate a splicing suppressor that results in skipping of exon 4, leading to frame shift and truncation of the protein (p. R181SfsX5) [31,32,34]. The c.626C site 20 base pairs (bp) upstream of the 3’ end of exon 4 is predicted to eliminate the two overlapping exonic splicing enhancers (ESE) (GT**C**AAG and T**C**AAGA) and generate an exonic splicing suppressor (ESS) sequence (GTTGT**T**) [35,36]. To confirm the expected exon skipping and truncation, we performed differential splicing analysis of RNA sequencing data obtained from the primary fibroblast cells from three patients carrying the homozygous c. 626 C>T mutation and five unaffected controls using Bisbee. We found that the MTFMT exon 4 skipping event was the 14th highest scoring differentially spliced event. Though the LLR (7.999) was just barely below the optimal threshold determined in the GTEx analysis, its high rank makes it likely to be considered in a candidate variant analysis. It is not surprising that the event did not quite pass the threshold as coverage of the event in the cases was only 10, 6 and 2 reads. If we use the protein effects predictions to filter down to events predicted to generate novel sequences that were expressed more highly in the cases compared to the controls, we find that the MTFMT exon 4 skip is the highest scoring of these events (Figure 5).

**Figure 5.**
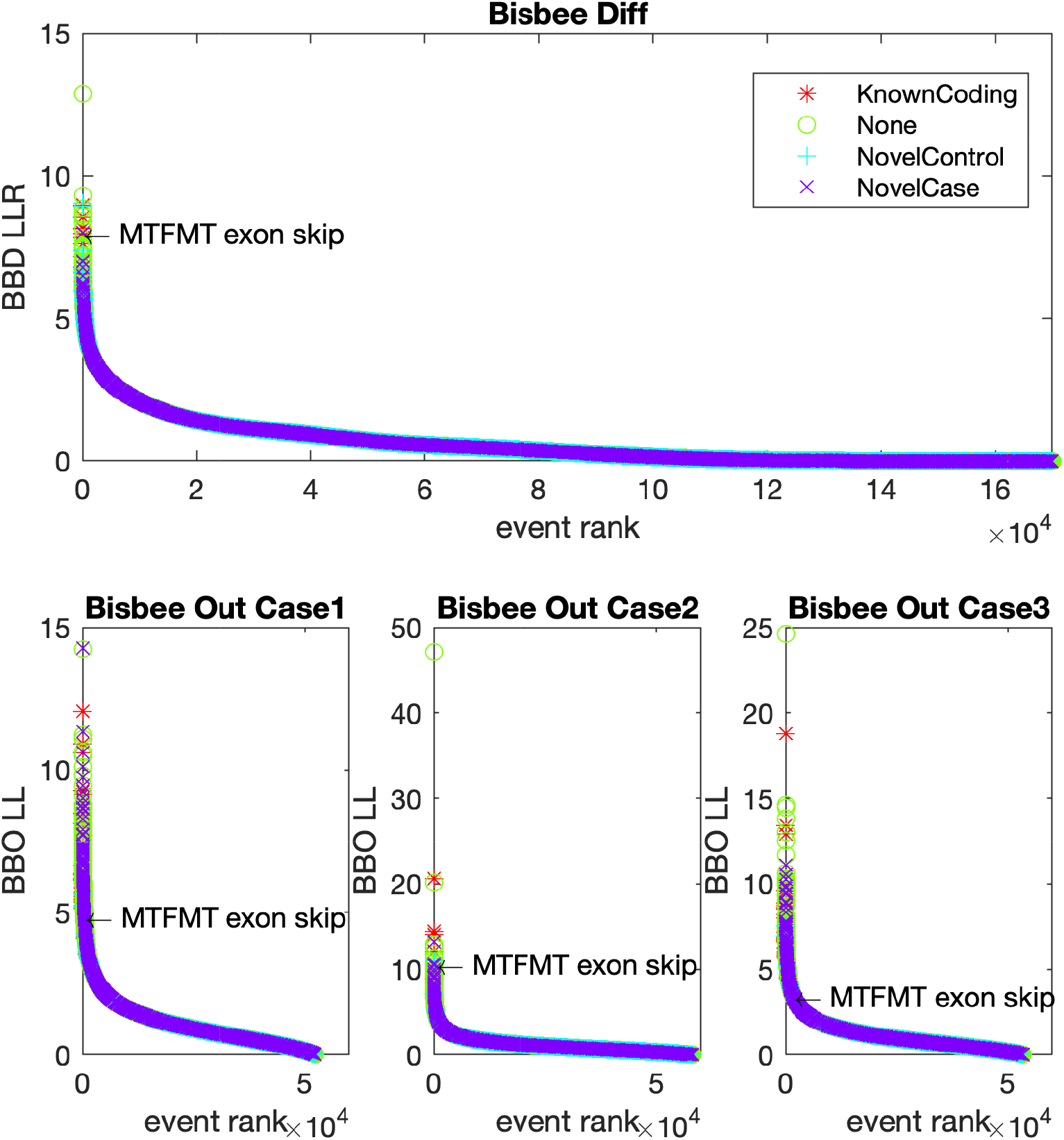
Bisbee detects known Leigh syndrome pathogenic splice variants in the MTFMT gene. The Bisbee scores are plotted on the y-axis against the rank of the scores on the x-axis in the differential splicing analysis (top) and the outlier analysis for the three cases (bottom). Each point represents a splice event and the color and symbol indicate whether the event is predicted to generate known coding isoforms (red asterisk), none coding or effect not predicted (green circle), novel coding sequence with the novel isoform more highly expressed in the controls (cyan plus) or more highly expressed in the cases (purple x).

When trying to discover the causal variant in a rare disease, there is often only one affected case available for sequencing, so we also ran Bisbee outlier analyses on each of the three cases to illustrate the single case scenario. Since it is desirable to have a large set of reference samples for outlier analysis, but there are technical differences in the sequencing between GTEx and this dataset, we performed the outlier analysis both using GTEx fibroblasts as the reference samples and using the five unaffected fibroblast samples used in the differential splicing analysis as the reference samples, and used the minimum score of the two analysis. The Bisbee outlier scores for the MTFMT exon 4 skip in the cases were 4.8, 10.6, and 3.4, ranking 386, 16, and 1746 of all events (Figure 5). When only considering events generating novel protein sequences with increased expression in the cases, the MTFMT event ranked 145, 2, and 587, respectively, in each of the three cases. Despite the very low coverage of the event in the cases, Bisbee was still able to rank the event in the top 1% of all events in all three cases.

The Bisbee annotation output is shown for the MFTMT exon 4 skipping event in Table 3. Each event is assigned a unique identifier (event_jid) using the contig, strand, and junction coordinates to facilitate comparing results between datasets. The effects at the transcript (event_cat) and protein level (effect_cat) are described, as well as whether the splice event is found in ensembl transcripts (aa_change_type). The sequence headers of the two isoforms are provided in order to locate the protein sequences in the fasta output. The sample group with increased expression of the isoform labeled “alt” is indicated (group_increased_alt). The location with the protein sequence as well as the altered amino acid sequence fragments are also provided. These results confirm the expected R181SfsX5 frame shift truncation.

**Table 3.**
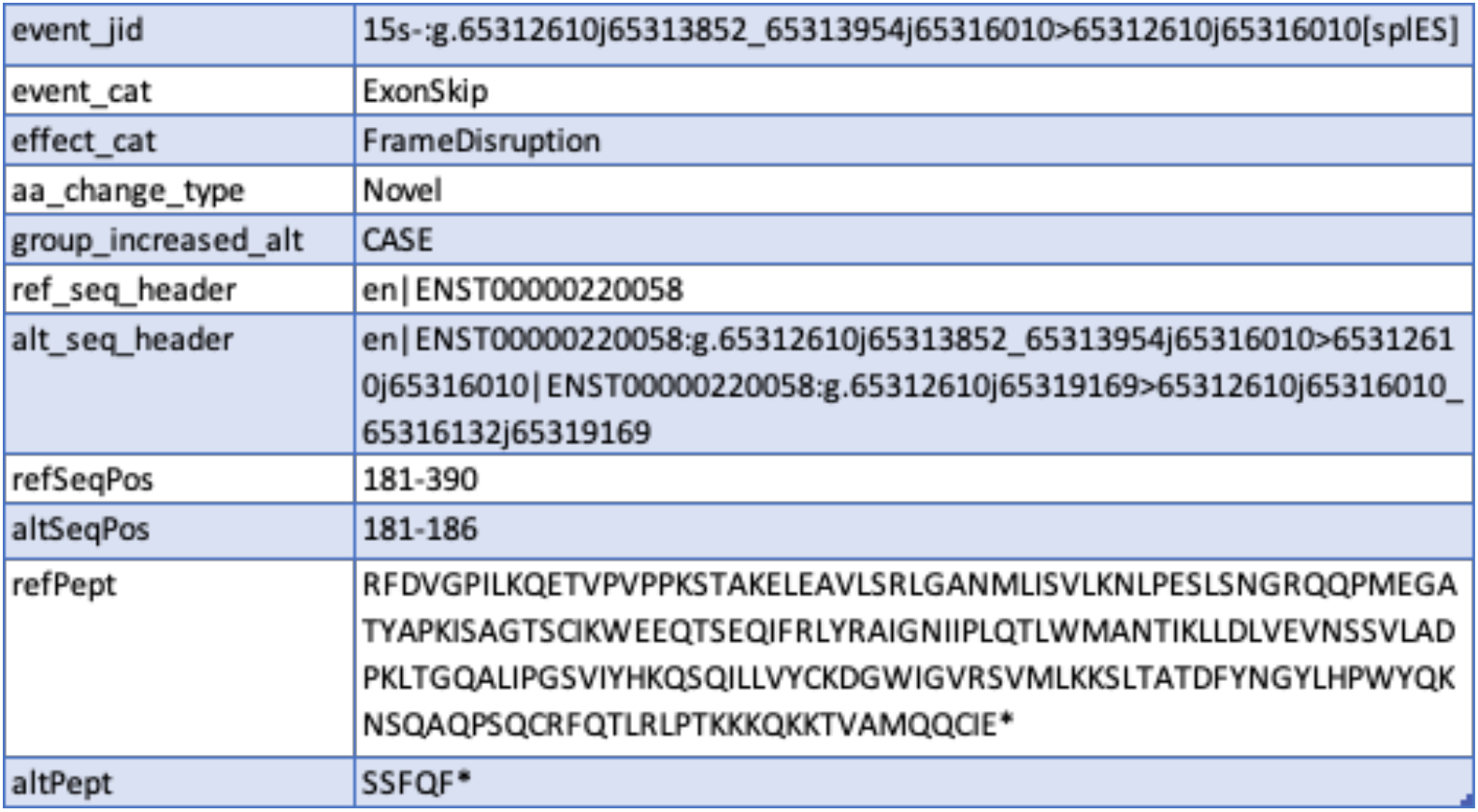
Example Bisbee output

### Application to TCGA Uveal Melanoma dataset

We selected the TCGA uveal melanoma dataset as an example application as there is a recurrent mutation in the splicing factor 3B1 gene (SF3B1) that has been previously shown to cause aberrant 3’ splice site usage [37,38]. To identify tumor-specific splice events, we performed Bisbee Outlier analysis using the complete GTEx tissue library exempt of testis tissue samples. Testis was excluded as it may express developmentally restricted proteins not found in normal somatic tissues [39,40]. We also used the TCGA normal samples as a reference and took the minimum score of the two analyses. In examining the total number of splice outliers per patient, we observed a large increase in alternative 3’ splice site outliers with SF3B1 mutation as well as significantly increased exon skipping, intron retention, and mutually exclusive exon outlier burden (Figure 6a, rank sum p-value <0.01). We also ran Bisbee Diff to identify differentially spliced events between SF3B1 mutant and wild-type tumors. We found 19,950 differentially spliced events of which 72% were mutually exclusive exons and 15% were alternative 3’ splice sites. The alternative 3’ differentially spliced events had higher Bisbee Diff LLR and a greater overlap with events also observed in the outlier analysis (Figure 6b). Alsafadi *et al.* previously identified differentially spliced events between SF3B1 mutant and wildtype tumors in an independent dataset, and selected seven of these events to validate in isogenic cell lines using a mini-gene splice assay [38]. All seven of these events were detected as differentially spliced by Bisbee Diff (Table 4).

**Figure 6.**
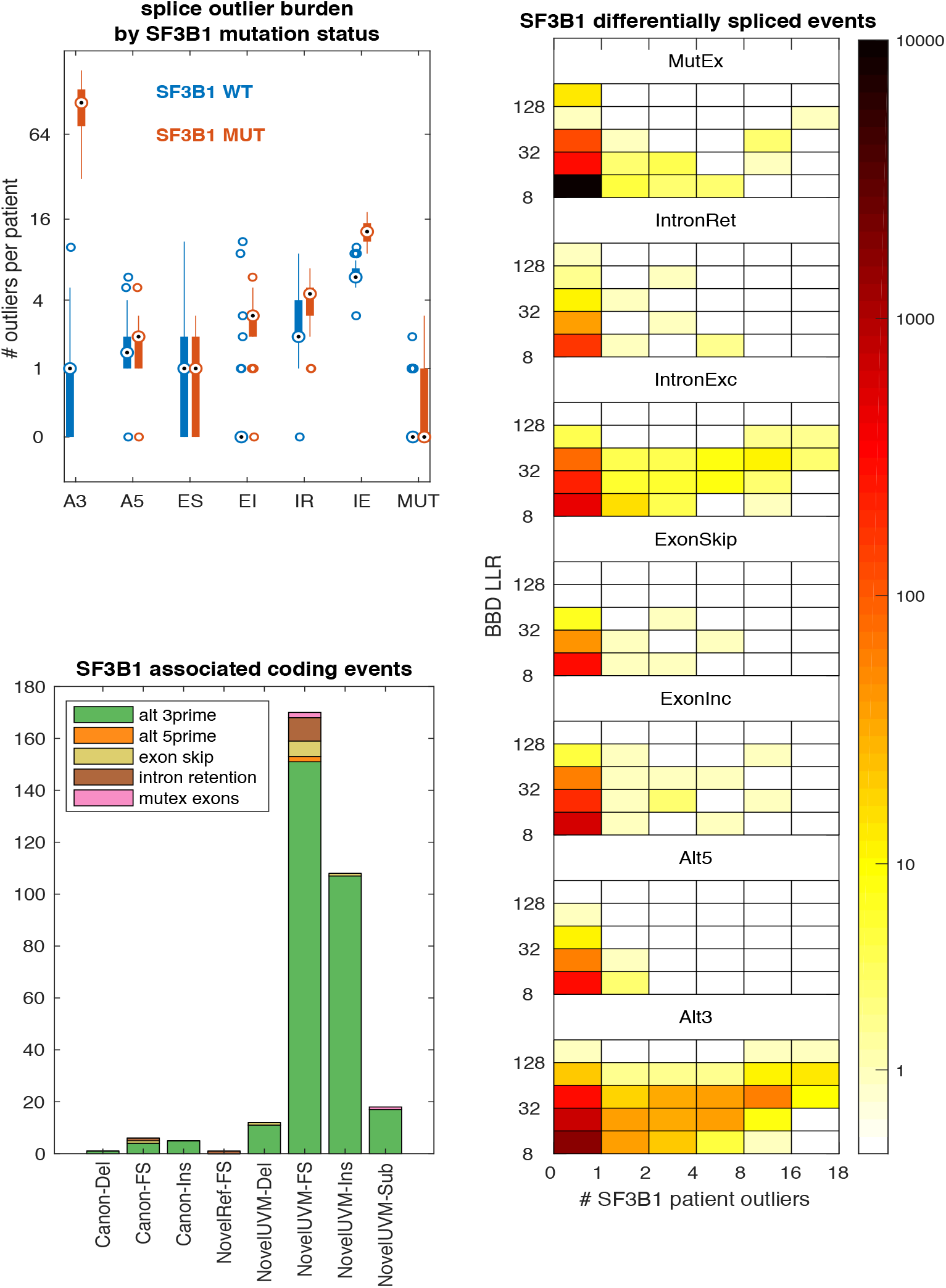
Aberrant splicing in uveal melanoma. a) Boxplot comparing the number of splice event outliers between SF3B1 mutant patients (n=18) and wild-type patients (n=62) by splice event type. b) Heatmap overview of differentially spliced events between SF3B1 mutant and wild-type tumors. Bins on the x-axis indicate the number of SF3B1 mutant tumors meeting the outlier criteria and bins on the y-axis indicate the Bisbee Diff LLR, and the color indicates the number of events falling into each bin. c) Event type and protein level effects of events that are differentially spliced between SF3B1 mutant and wildtype tumors, are outliers in at least one SF3B1 mutant tumor, and result in an altered protein sequence.

**Table 4.** Validated SF3B1 mutant vs wild-type differentially spliced events

| event jid | gene | effect cat | BBD<br>LLR | BBD<br>rank | fit PSI<br>SF3B1 | fit PSI<br>wt | # SF3B1 mutant<br>tumor outliers |
| --- | --- | --- | --- | --- | --- | --- | --- |
| 6s+:g.10723474j10724789><br>10723474j10724803[splA3] | TMEM14C | Silent | 148.9 | 16 | 0.76 | 0.03 | 18 |
| 1s-:g.101458310j101460666><br>101458296j101460666[splA3] | DPH5 | Frame<br>Disruption | 87.5 | 33 | 0.78 | 0.00 | 18 |
| 2s+:g.232196609j232209661><br>232196609j232209687[splA3] | ARMC9 | Frame<br>Disruption | 80.2 | 41 | 0.66 | 0.00 | 18 |
| 14s+:g.75356052j75356581><br>75356052j75356600[splA3] | DLST | Frame<br>Disruption | 76.6 | 44 | 0.50 | 0.01 | 0 |
| 18s-:g.683380j685921><br>683395j685921[splA3] | ENOSF1 | Insertion | 74.2 | 47 | 0.71 | 0.03 | 16 |
| 6s+:g.35255622j35258030><br>35255622j35258043[splA3] | ZNF76 | Frame<br>Disruption | 53.8 | 194 | 0.44 | 0.03 | 11 |
| 16s+:g.843274j844034><br>843274j844054[splA3] | CHTF18 | Frame<br>Disruption | 25.1 | 1316 | 0.30 | 0.00 | 8 |

In order to identify protein isoforms that may be specific to SF3B1 mutant tumors, we selected splice events that were common between the differential splicing and outlier analysis (494) and then identified those predicted to result in altered protein sequence (321). These events are primarily alternative 3’ events causing insertions or frame disruptions resulting in novel protein isoforms in the uveal melanoma tumors (Figure 6c).

### Replication of common melanoma associated splice events in an independent dataset

In addition to observing splice events associated with SF3B1 mutation, we also observed splice events common across the TCGA uveal melanoma cohort, irrespective of SF3B1 mutation status. In order to validate this finding, we performed the Bisbee splicing analysis on an independent melanoma cohort, consisting of 37 patients with BRAF wild-type recurrent tumors including 13 cutaneous, 7 mucosal, 10 uveal, 5 acral, 1 melanoma of unknown primary. We performed the Bisbee outlier analysis using both the GTEx excluding testis as the reference and a set of 28 normal tissue or cell lines sequenced at the same institution as the reference and took the minimum score of the two analyses. We compared the number of patients passing the outlier threshold for each event between the two datasets. We identified 23 splice events with 20 or more tumors meeting the outlier criteria in the TCGA dataset, and found that 10 of these events were also detected as outliers in at least one of the SU2C tumors (Figure 7a). When only considering events with predicted protein sequence changes, there are ten events meeting the outlier criteria in 20 or more of the TCGA tumors and nine of these events are detected as outliers in at least one of the SU2C tumors (Figure 7b). These nine events identified in both datasets include five intron exclusion events in GAPDHS predicted to result in novel sequence in the reference samples. There is also an alternative 5 prime site in EXOC3, and intron retention in TBL1X, PTPRH, and PALM that are predicted to result in novel sequence in the tumors (Figure 7c). The intron retention event in SLC24A5 was not detected by SplAdder in the SU2C dataset.

**Figure 7.**
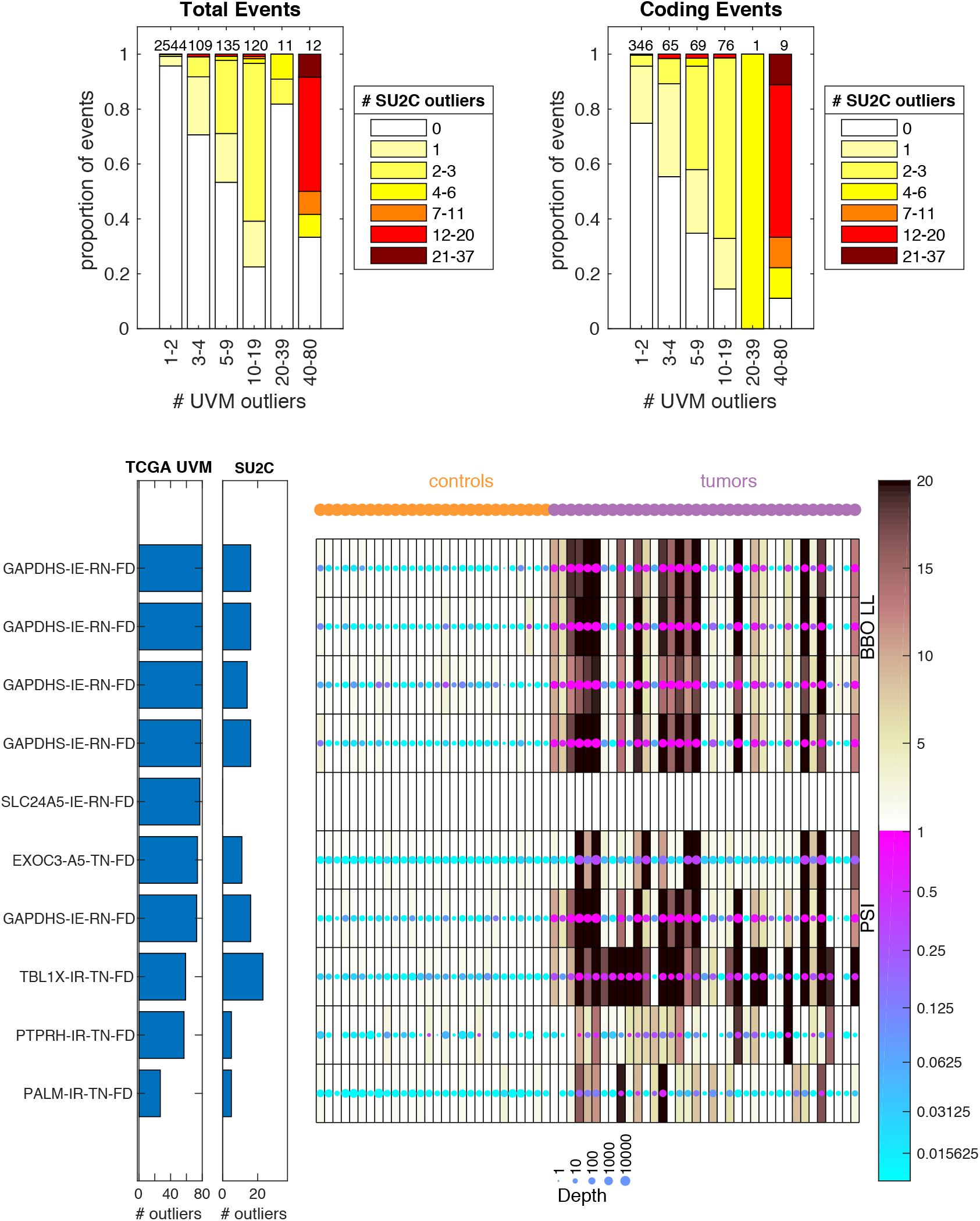
Common melanoma associated splice events are shared between cohorts. a) Comparison of number tumors meeting outlier criteria between cohorts. Along the x-axis, events are binned by the number of tumors meeting the outlier threshold in the TCGA cohort. The total number of events in each bin is indicated at the top of each bar. Within each bar, the events are binned by the total number of tumors meeting the outlier threshold in the SU2C cohort, and the proportion of events in each bin is indicated by the color. b) Same as a) but only including events with predicted protein coding changes. c) Heatmap of the data in the SU2C cohort for the 10 events with predicted coding sequence changes that are found in more than 20 tumors in the TCGA cohort. Each row is an event and each column is a sample, with the controls on the left and the tumors on the right. The color of each dot represents the PSI and the size of each dot represents the coverage at the event. The shading behind the dot indicates the Bisbee outlier score. The bar graphs to the left of the heatmap indicate the number of tumors meeting the outlier threshold in each cohort. The labels on the left indicate the gene name, event type (IE – intron exclusion, A5 – alternative 5’, IR – intron retention), and the effect type (RN-FD – frame disruption in the control samples predicted to result in novel sequences, TN-FD frame disruption in the tumor samples predicted to result in novel sequences).

## Discussion

We have developed a new package for splicing data analysis called Bisbee. Bisbee provides functions for differential splicing analysis, splicing outlier analysis, and protein effect prediction. Using a dataset with matched RNAseq and mass spectrometry data on normal human tissues we constructed a truth set to benchmark differential splicing and outlier methods including Bisbee. We found that Bisbee’s differential splicing approach had substantially better enrichment of proteomics confirmed events than the other approaches. Bisbee’s outlier test also outperformed other outlier approaches. We demonstrated the utility of the approach in both a rare disease and a cancer context.

The Bisbee package goes beyond many other splicing tools by generating protein sequences for the observed splice events. In addition to enabling proteomics-based benchmarking, the generation of protein sequences also enable a more integrated analysis of patient samples including mass spectrometry based approaches for detection of splicing-derived proteoforms. In order to identify protein isoforms, a database of protein sequences is required to match against mass spectra. By using Bisbee to generate a database of patient specific protein isoforms from RNAseq data, and then using mass spectrometry to detect which ones have protein level evidence, one could identify high confidence disease-specific protein isoforms for further characterization. The protein domains impacted and other downstream functional predictions from the protein sequences enable further insight into the impact of splicing alterations and can identify splicing-derived pathogenic variants that would go undetected by DNA sequencing alone.

The Bisbee pipeline currently relies on SplAdder for splice event detection [14]. While the work presented here as well as previous work demonstrate that SplAdder is a robust tool for splice event detection, in the future, we plan to adapt Bisbee to work with input from other splice detection tools and benchmark against SplAdder. Bisbee is also limited to the types of splice events detected by SplAdder: alternate 3’ and 5’ splice sites, exon skipping, intron retention and mutually exclusive exons. SplAdder recently added multiple exon skips detection but that is not yet supported by Bisbee and GTEx results are not available for this event type. Other event types that are not currently detected include alternate first exon, alternate terminal exon, and complex events involving more than one type of alteration. Another limitation of the current approach is that it relies on short read sequencing and does not attempt to assemble a fulllength transcript but rather focuses on the local changes in the transcript and protein sequences. An expansion of the approach to incorporate long read data would be useful for enabling full length sequence analysis. Currently, Bisbee only offers two statistical tests: comparison between two groups and outlier detection compared to a reference set. Future work may extend the methods to test for associations with continuous variables or other more complex experimental designs.

The differential splicing test in Bisbee uses a novel beta binomial model to test for differences in PSI. Most differential splicing tools, including the SplAdder test included in our evaluation, test for differences in expression level of the splice isoform, controlling for the overall expression level of the gene. Many of the events that are highly significant in SplAdder’s test have relatively small differences in mean PSI between the two groups (Figure 4b). In order to identify events with more substantial differences in mean PSI between the two groups, one may directly test for a difference in PSI values using a t-test. However, this does not take into account that PSI measurements are inherently more noisy for events with lower coverage. While using a depth threshold on the PSI measurements considered for differential expression will eliminate some of the false positives due to noisy PSI measurements, we have shown that the beta binomial model implemented in Bisbee better addresses the relationship between PSI measurement accuracy and depth. Bisbee is able to detect both low coverage events with dramatic differences in PSI and high coverage events with small differences in PSI (Figure 4a).

Bisbee is the second splicing tool that we are aware of to offer an outlier detection test. This test is intended for identifying splice isoforms unique to an individual patient compared to a set of reference samples. We were not able to compare directly to the other splice outlier detection tool (leafCutterMD) as it does not report splice events in a way that is amenable to protein sequence generation. Further application of this tool may enable the discovery of splicing defects that are causal in rare genetic disorders or specific to an individual patient’s tumor. The case studies we presented illustrate the utility of the outlier approach in both the rare disease and cancer research.

We also discussed an example analysis demonstrating the detection of a known splice event due to a pathogenic variant in a rare disease. Since three cases with known pathogenic splice variants were available, we were able to perform differential splicing analysis. The casual event was the 14th highest scoring of all differentially spliced events, and the highest scoring of those predicted to generate a novel amino acid sequence, illustrating how the protein level annotation can aid variant prioritization. Outlier analysis is an important approach in studying rare disease as often more than one case is not available. These cases were difficult to detect by the outlier analysis due to the very low coverage at the event locus. One of the three cases had the casual event ranked 33rd overall and 2nd of events generating novel protein sequences in spite of having a coverage of only 6 reads at the event locus. While the event was not ranked as highly in the other two cases, it is conceivable that the Bisbee output could still help identify the causal variant when examined along with occurrence of rare variants in the region and knowledge of the phenotype and underlying pathways is exploited.

Previous work has suggested that splicing dysregulation in cancer may be a greater source of tumor specific antigens than somatic point mutations [3,25]. Application of the Bisbee outlier test to cancer patient samples may enable the discovery of tumor-specific splicing-derived neoantigens, which could be therapeutic or vaccine targets. In our reanalysis of the TCGA uveal melanoma dataset, we found that Bisbee robustly detects the independently validated aberrant 3’ splice site usage associated with SF3B1 mutation [37,38]. Splice events that are both outliers compared to normal tissues and differentially spliced between SF3B1 mutant and wild-type tumors are promising candidates as tumor-specific neoantigens, as many of these are predicted to generate novel sequences through frame disruptions and insertions in the tumor-specific isoforms (Figure 6c). SF3B1 mutant uveal melanomas have better prognosis than SF3B1 wild-type [39]. We hypothesize that the tumor-specific splice isoforms associated with SF3B1 mutations may act as antigens enabling better immune control of the tumors. The protein sequence output from Bisbee would facilitate in silico MHC binding prediction to further investigate the potential immunogenicity of these splice variant generated neoantigens.

We also detected splice outliers common to uveal melanoma regardless of SF3B1 mutation status, and these results showed strong concordance in an independent melanoma cohort. Interestingly, events with predicted protein sequence impact showed stronger concordance than those with no predicted impact (Figure 7a and b). Nine of the ten events identified as common splice variant outliers with protein impact in the TCGA uveal melanoma dataset were also detected in the SU2C melanoma dataset. These melanoma associated splice variants included several intron retention events in GAPDHS, with the tumors having lower expression of the intron-retained transcripts compared to the normal reference tissues. GAPDHS is typically expressed in sperm, but not in normal somatic tissues, and has previously been shown to be expressed in melanoma [40]. We hypothesize that we are seeing these events in GAPDHS due to expression of the immature transcript in the normal tissues. Four melanoma associated splice events were identified that were predicted to lead to frame disruptions in the tumors, resulting in novel protein sequence. These events are most promising for further investigation as candidate targets in melanoma.

## Conclusions

The Bisbee package is able to predict protein sequences of both known and novel protein isoforms. It provides a more statistically powerful differential splicing test than existing methods. It also provides an outlier detection approach, which will be useful in a number of different contexts, including cancer and rare disease. We presented an example analysis of a rare disease where Bisbee was able to highly rank the splice event resulting from the pathogenic variant in spite of very low read coverage of the event. We also applied the Bisbee package to the TCGA uveal melanoma dataset and were able to robustly detect the expected association between SF3B1 mutation and aberrant 3’ splice site usage. We were also able to detect the common splice outliers across uveal melanoma tumors that were replicated in an independent dataset. The Bisbee package is publicly available, and should enable the robust detection of aberrant splicing.

## Methods

### Description of datasets used

For initial evaluation and optimization of the differential and outlier splicing test implemented in Bisbee, we compared the distribution of the likelihood ratios between tests involving samples from the same tissues compared to samples from different tissues. We would expect that different tissues should have true differences in splicing events, so greater enrichment for higher scores in the different tissues comparison would indicate a better powered test. For this analysis GTEx SplAdder results were downloaded from GDC (https://gdc.cancer.gov/about-data/publications/PanCanAtlas-Splicing-2018) [3]. For the differential splicing evaluation, 50 random pairs of tissues were selected, six random samples were selected from each tissue, and 100,000 events were selected for each tissue pair. The beta binomial differential splicing test was applied to grouping the samples into two groups of three replicates within each tissue as well as between the pairs of different tissues. For the outlier evaluation, 12 tissues with at least 100 samples were selected. For each of these 12 tissues, 80 samples were randomly selected for fitting the model and 20 were selected for determining the outlier scores.

For further evaluation and benchmarking, we identified a dataset where RNAseq and mass spectrometry data were available on the same set of tissues [8]. RNAseq reads were downloaded from ArrayExpress (E-MTAB-2836) and aligned to the human reference genome (GRCh38) using star 2.7.3a two pass basic mapping mode and splice events were detected using SplAdder. A concatenated fasta containing protein sequences predicted for all splicing events and canonical sequences from Ensembl was used as the search database for LC-MS/MS spectra downloaded for 7 tissue types from the EBI PRIDE database (PXD010154). The spectra were searched using Mascot (Matrix Science, London, UK; version 2.6.0) through Proteome Discoverer 2.4 (Thermo Fisher Scientific, Waltham, MA), allowing for oxidation (Met) and carbamidomethylation (Cys) dynamic and static modifications, respectively. A maximum of two missed cleavages were allowed with fragment mass tolerance of 0.02Da and precursor mass tolerance of 10ppm. FDR thresholds for PSMs, peptides and proteins were set at 0.01, with a minimum of 1 peptide required for protein identification. Peptides that matched exclusively to only one protein sequence were taken as evidence for that isoform. Events where only one isoform was detected in one tissue and the other isoform detected in a different tissue were taken as protein-level evidence of tissue-specific splicing.

For an example use case, three Leigh syndrome and five unaffected control fibroblast cell lines from the study participants were established from 3 mm skin biopsy punches and cultured for 2 weeks in primary fibroblast media containing the following: Minimal Essential Media (Invitrogen, Carlsbad, CA, USA), 20% FBS (American Type Culture Collection, Manassas, VA, USA), Penicillin/Streptomycin and Amphotericin (Sigma-Aldrich, St. Louis, MO, USA), and Plasmocin (InvivoGen, San Diego, CA, USA) (Villegas & McPhaul 2005). RNA was extracted from the study fibroblast using the total RNA Purification Kit (Norgen Biotek Corp, On, Canada), and RNA Library Preparation was done with the KAPA mRNA HyperPrep Kit with RiboErase (Kapa Biosystems) according to the manufacturer’s protocol. Barcoded libraries were pooled (8-plex) and run on two lanes of a flow cell on an Illumina NovaSeq 6000 System. Sequencing was performed using 2 × 100 bp reads. The dataset was aligned to the human reference genome GRCh37 using STAR (v2.4.0) [41], and quality control was performed using Picard RnaSeqMetrics (v1.128).

For the uveal melanoma analysis, TCGA SplAdder results were downloaded from GDC (https://gdc.cancer.gov/about-data/publications/PanCanAtlas-Splicing-2018) [3]. The SF3B1 mutation status was obtained from cBioportal (https://bit.ly/3hagZvp) [42].

We used an independent set of melanoma patients for comparison with the TCGA melanoma dataset, referred to here as the SU2C melanoma cohort. Biopsies were collected as part of a clinical trial (NCT02094872), where inclusion criteria included patients aged ≥18 years with metastatic or locally advanced and unresectable *BRAFwt* melanoma who had progressed following previous treatment. RNA was extracted from frozen core needle biopsies using the Qiagen AllPrep Kit. Illumina’s TruSeq RNA Sample Preparation V2 Kit was used to construct libraries which were sequenced on the Illumina HiSeq2500 for 2×100 reads. RNA FASTQs were aligned to the human reference genome GRCh37using STAR 2.3.1z [41].

### Splice event detection

We used SplAdder v2.3.0 for splice event detection with default parameters. The coordinates of the junctions involved in each event, and the counts of reads supporting isoform 1 and isoform 2 for each event in each sample were extracted from the counts hdf5 files.

### Splice event protein sequence prediction

In order to generate protein sequences corresponding to each splice event, we use known transcript sequences from Ensembl as a starting point. We first determined whether each isoform of the event exists with any known transcripts. We compared the event junction coordinates to the exon coordinates of protein coding transcripts for that gene using the python package pyEnsembl to retrieve the exon coordinates. Each transcript is categorized as matching isoform one, isoform two, or neither for the splice event. For each transcript matching the isoform one, the isoform one junctions are removed and replaced with the isoform two junctions to make the altered sequence, and vice versa for those matching isoform two. The region of altered amino acids is found by aligning the two sequences. If the altered amino acid sequence is not found in any of the canonical sequences, the event is categorized as novel. If no transcript is found that matches either isoform, no sequence is generated and the event’s effect is categorized as unknown. In order to narrow down to one pair of protein sequences per event, the sequences are prioritized as follows: 1) pair of known transcripts 2) longest altered amino acid sequence 3) longest starting isoform sequence.

### Differential splicing test (Bisbee diff)

Read counts for a splice variant are modeled as following a beta binomial distribution, which is similar to a binomial distribution in that it models the number of successes given a fixed number of trials, but rather than having a constant probability of success, the probability of success follows a beta distribution. Here the number of reads supporting the first isoform is the number of successes, the total number of reads covering the event is the number of trials, and the expected distribution PSI (percent spliced in) across the samples of interest is represented by the beta distribution. The beta distribution is reparameterized as 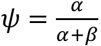 and *ω* = *α* + *β* so that *ψ* represents the expected value of the beta distribution and *ω* affects the sharpness of the distribution. In the one group model, all of the samples are assumed to have the same underlying distribution of PSI values and a maximum likelihood estimate is made for *ψ* and *ω*.

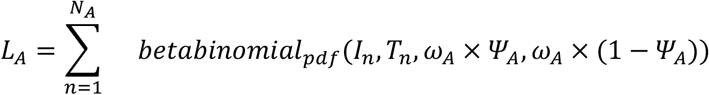

In the two-group model, it is assumed that the two groups have different expected PSI values, but similar distribution shapes, so the two groups have different values of *ψ* but the same *ω*.

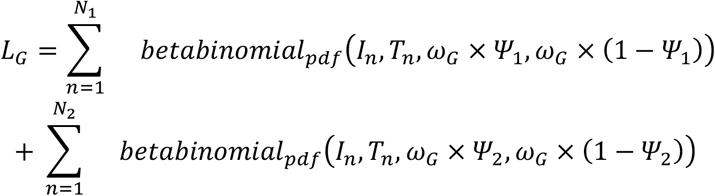

For both the one group and two group models the values of the parameters that maximize the sum of the probability densities across the data points. In fitting the model, we use the following transformations to constrain ωto be greater than 2 and less than *ω_M_* and constrain *Ψ* to be between 0 and 1.

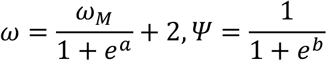

The difference in the sum of the log probability densities across the two models is used to identify that events have different underlying PSI distributions in the two groups.

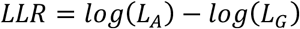

### Splicing outlier test (Bisbee outlier)

As in the two-group test, the read counts are assumed to follow a beta binomial distribution, where the number of reads supporting the first isoform is considered the number of successes, the total coverage at the locus is the number of trials, and the expected distribution of PSI values is represented by the beta distribution. The beta distribution parameters are found that maximizes the sum of the log probability densities across a set of reference samples.

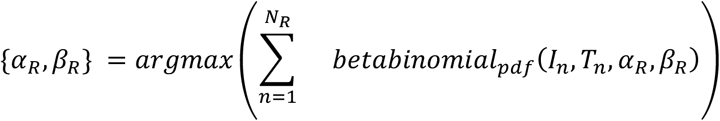

Here *I_n_* indicates the read count supporting the isoform with mean PSI<0.5 across the samples. Nelder-Mead optimization [44] (or BFGS if Nelder-Mead fails) is used to find the maximum likelihood values of *α_R_,β_R_*. The reparameterizations below are used to constrain *α_R_* to be between 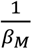 and 1 and *β_R_* to be between 1 and *β_M_*so that the beta distribution is strictly decreasing.

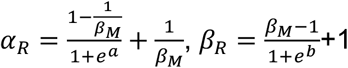

If zero reads are detected supporting the minor isoform in the reference sample set, alpha is set to one and beta is set as shown below.

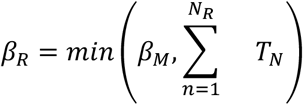

For each sample of interest, the log cumulative probability of the major isoform read counts being less than or equal to those observed given the total read depth and the beta distribution fit to the reference sample is used as a high outlier score.

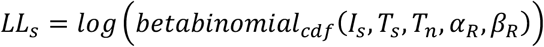

### Implementation of other differential splicing methods

For benchmarking we selected SplAdder’s differential splicing test using default parameters as a representative of the generalized linear model approach. We also wanted to include a simple method directly testing differences in PSI values. While a non-parametric test would be more appropriate, as PSI values are unlikely to be normally distributed, we would not have any power to detect differences with only three or four replicates per group. Instead, used a two-sample t-test, with the more conservative assumption of unequal variance, on the PSI values. SplAdder only reports PSI values for samples with a coverage of 10 for a given event, though PSI values can still be calculated from the isoform one and two coverages. We applied the t-test both to all PSI values as well as treating the data points with depth less than 10 as missing data.

### Implementation of other splicing outlier detection methods

As we are not aware of other tools to detect splice outliers, we implemented two simple methods using the distribution of PSIs for comparison. The first finds the median absolute deviation from the set of reference samples.

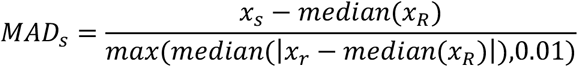

The second normalizes to the interquartile range.

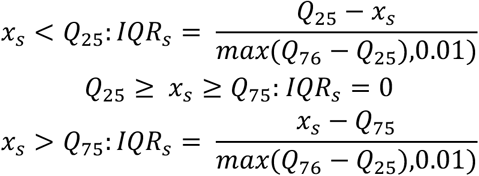

We applied both of these methods either using all of the PSI values, as well as only using data points with depth less than 10 as missing data.

## Declarations

### Ethics approval and consent to participate

The study protocol and written informed consent for the use of human fibroblast cells of the MFTMT cases and controls was approved by the Western Institutional Review Board (WIRB; study number 20120789). The SU2C melanoma biopsies were collected through a clinical trial (NCT02094872). Ethics review boards at all participating institutions approved the study, which was conducted in accordance with the Declaration of Helsinki and Good Clinical Practice guidelines. All patients provided written informed consent.

### Consent for publication

Not applicable

### Availability of data and materials

The RNA sequencing data of the MTFMT cases and controls will be deposited in dbGap. The SU2C RNA sequencing data is available in dbGap under accession phs001786.v1.p1. The Bisbee package is available at https://github.com/tgen/bisbee

### Author Contributions

RFH designed and implemented the Bisbee package and performed the splicing analysis. AH performed the proteomics analysis and beta tested the Bisbee package. JDL, EAR, PP, and NJS contributed to the interpretation of the data and writing of the manuscript. RS, CL, and the C4RCD research group designed and performed the experiments and contributed to the analysis and interpretation of the rare disease dataset. WSL, PML, AS, JAS, and JMT designed and performed the experiments and contributed to the analysis and interpretation of the melanoma dataset. PP conceived the proteomics evaluation strategy and NJS contributed to the statistical analysis. All authors have read and approved the final manuscript.

## Acknowledgements

The authors would like to thank Krystine Garcia-Mansfield for input on setting up the mass spectrometry searches and Megan Johnson for beta testing and debugging the Bisbee package. The C4RCD research group includes Vinodh Narayanan, Matt Huentelman, Newell Belnap, Anne-Marie Aziz, and Keri Ramsey.

## References

1. Park E, Pan Z, Zhang Z, Lin L, Xing Y. The Expanding Landscape of Alternative Splicing Variation in Human Populations. Am J Hum Genet. 2018;102:11–26.

2. Gamazon ER, Stranger BE. Genomics of alternative splicing: evolution, development and pathophysiology. Hum Genet. 2014;133:679–87.

3. Kahles A, Lehmann K-V, Toussaint NC, Hüser M, Stark SG, Sachsenberg T, et al. Comprehensive Analysis of Alternative Splicing Across Tumors from 8,705 Patients. Cancer Cell. 2018;34:211–224.e6.

4. Dvinge H, Kim E, Abdel-Wahab O, Bradley RK. RNA splicing factors as oncoproteins and tumor suppressors. Nat Rev Cancer. 2016;16:413–30.

5. Li YI, Geijn B van de, Raj A, Knowles DA, Petti AA, Golan D, et al. RNA splicing is a primary link between genetic variation and disease. Science. 2016;352:600–4.

6. Barbeira AN, Bonazzola R, Gamazon ER, Liang Y, Park Y, Kim-Hellmuth S, et al. Widespread dose-dependent effects of RNA expression and splicing on complex diseases and traits. bioRxiv. 2019;814350.

7. Anna A, Monika G. Splicing mutations in human genetic disorders: examples, detection, and confirmation. J Appl Genet. 2018;59:253–68.

8. Wang D, Eraslan B, Wieland T, Hallström B, Hopf T, Zolg DP, et al. A deep proteome and transcriptome abundance atlas of 29 healthy human tissues. Mol Syst Biol. 2019;15:e8503.

9. Frazee AC, Pertea G, Jaffe AE, Langmead B, Salzberg SL, Leek JT. Ballgown bridges the gap between transcriptome assembly and expression analysis. Nat Biotechnol. 2015;33:243–6.

10. Katz Y, Wang ET, Airoldi EM, Burge CB. Analysis and design of RNA sequencing experiments for identifying isoform regulation. Nat Methods. 2010;7:1009–15.

11. Shen S, Park JW, Lu Z, Lin L, Henry MD, Wu YN, et al. rMATS: Robust and flexible detection of differential alternative splicing from replicate RNA-Seq data. Proc Natl Acad Sci. 2014;111:E5593–601.

12. Trincado JL, Entizne JC, Hysenaj G, Singh B, Skalic M, Elliott DJ, et al. SUPPA2: fast, accurate, and uncertainty-aware differential splicing analysis across multiple conditions. Genome Biol [Internet]. 2018 [cited 2018 Aug 13];19. Available from: https://www.ncbi.nlm.nih.gov/pmc/articles/PMC5866513/

13. Mancini E, Iserte J, Yanovsky M. ASpli: An integrative R package for analysing alternative splicing using RNA-Seq.:34.

14. Kahles A, Ong CS, Zhong Y, Rätsch G. SplAdder: identification, quantification and testing of alternative splicing events from RNA-Seq data. Bioinformatics. 2016;32:1840–7.

15. Goldstein LD, Cao Y, Pau G, Lawrence M, Wu TD, Seshagiri S, et al. Prediction and Quantification of Splice Events from RNA-Seq Data. PLoS ONE [Internet]. 2016 [cited 2017 Aug 30];11. Available from: http://www.ncbi.nlm.nih.gov/pmc/articles/PMC4878813/

16. Li YI, Knowles DA, Humphrey J, Barbeira AN, Dickinson SP, Im HK, et al. Annotation-free quantification of RNA splicing using LeafCutter. Nat Genet. 2018;50:151–8.

17. Vaquero-Garcia J, Barrera A, Gazzara MR, González-Vallinas J, Lahens NF, Hogenesch JB, et al. A new view of transcriptome complexity and regulation through the lens of local splicing variations. Valcárcel J, editor. eLife. 2016;5:e11752.

18. Gerstung M, Beisel C, Rechsteiner M, Wild P, Schraml P, Moch H, et al. Reliable detection of subclonal single-nucleotide variants in tumour cell populations. Nat Commun. Nature Publishing Group; 2012;3:811.

19. Christoforides A, Carpten JD, Weiss GJ, Demeure MJ, Hoff DDV, Craig DW. Identification of somatic mutations in cancer through Bayesian-based analysis of sequenced genome pairs. BMC Genomics. 2013;14:302.

20. Halperin RF, Liang WS, Kulkarni S, Tassone EE, Adkins J, Enriquez D, et al. Leveraging spatial variation in tumor purity for improved somatic variant calling of archival tumor only samples. Front Oncol [Internet]. 2019 [cited 2019 Feb 14];9. Available from: https://www.frontiersin.org/articles/10.3389/fonc.2019.00119/abstract

21. Shiraishi Y, Sato Y, Chiba K, Okuno Y, Nagata Y, Yoshida K, et al. An empirical Bayesian framework for somatic mutation detection from cancer genome sequencing data. Nucleic Acids Res. 2013;41:e89.

22. Urbanski LM, Leclair N, Anczuków O. Alternative-splicing defects in cancer: Splicing regulators and their downstream targets, guiding the way to novel cancer therapeutics. Wiley Interdiscip Rev RNA. 9:e1476.

23. Jayasinghe RG, Cao S, Gao Q, Wendl MC, Vo NS, Reynolds SM, et al. Systematic Analysis of Splice-Site-Creating Mutations in Cancer. Cell Rep. 2018;23:270–281.e3.

24. Hoyos LE, Abdel-Wahab O. Cancer-Specific Splicing Changes and the Potential for Splicing-Derived Neoantigens. Cancer Cell. 2018;34:181–3.

25. Frankiw L, Baltimore D, Li G. Alternative mRNA splicing in cancer immunotherapy. Nat Rev Immunol. 2019;19:675–87.

26. Jenkinson G, Li YI, Basu S, Cousin MA, Oliver GR, Klee EW. LeafCutterMD: an algorithm for outlier splicing detection in rare diseases. Bioinformatics [Internet]. [cited 2020 May 1]; Available from: https://academic.oup.com/bioinformatics/advance-article/doi/10.1093/bioinformatics/btaa259/5823301

27. Boise LH, González-García M, Postema CE, Ding L, Lindsten T, Turka LA, et al. bcl-x, a bcl-2-related gene that functions as a dominant regulator of apoptotic cell death. Cell. Elsevier; 1993;74:597–608.

28. Zhang H, Liu T, Zhang Z, Payne SH, Zhang B, McDermott JE, et al. Integrated Proteogenomic Characterization of Human High-Grade Serous Ovarian Cancer. Cell. 2016;166:755–65.

29. Mertins P, Mani DR, Ruggles KV, Gillette MA, Clauser KR, Wang P, et al. Proteogenomics connects somatic mutations to signalling in breast cancer. Nature. 2016;534:55–62.

30. Vasaikar S, Huang C, Wang X, Petyuk VA, Savage SR, Wen B, et al. Proteogenomic Analysis of Human Colon Cancer Reveals New Therapeutic Opportunities. Cell. Elsevier; 2019;177:1035–1049.e19.

31. Tucker EJ, Hershman SG, Köhrer C, Belcher-Timme CA, Patel J, Goldberger OA, et al. Mutations in MTFMT Underlie a Human Disorder of Formylation Causing Impaired Mitochondrial Translation. Cell Metab. 2011;14:428–34.

32. Haack TB, Gorza M, Danhauser K, Mayr JA, Haberberger B, Wieland T, et al. Phenotypic spectrum of eleven patients and five novel MTFMT mutations identified by exome sequencing and candidate gene screening. Mol Genet Metab. 2014;111:342–52.

33. Hayhurst H, Coo IFM de, Piekutowska-Abramczuk D, Alston CL, Sharma S, Thompson K, et al. Leigh syndrome caused by mutations in MTFMT is associated with a better prognosis. Ann Clin Transl Neurol. 2019;6:515–24.

34. Neeve VCM, Pyle A, Boczonadi V, Gomez-Duran A, Griffin H, Santibanez-Koref M, et al. Clinical and functional characterisation of the combined respiratory chain defect in two sisters due to autosomal recessive mutations in MTFMT. Mitochondrion. 2013;13:743–8.

35. Fairbrother WG, Yeh R-F, Sharp PA, Burge CB. Predictive identification of exonic splicing enhancers in human genes. Science. 2002;297:1007–13.

36. Wang Z, Rolish ME, Yeo G, Tung V, Mawson M, Burge CB. Systematic identification and analysis of exonic splicing silencers. Cell. 2004;119:831–45.

37. Furney SJ, Pedersen M, Gentien D, Dumont AG, Rapinat A, Desjardins L, et al. SF3B1 Mutations Are Associated with Alternative Splicing in Uveal Melanoma. Cancer Discov. American Association for Cancer Research; 2013;3:1122–9.

38. Alsafadi S, Houy A, Battistella A, Popova T, Wassef M, Henry E, et al. Cancer-associated SF3B1 mutations affect alternative splicing by promoting alternative branchpoint usage. Nat Commun. Nature Publishing Group; 2016;7:10615.

39. Harbour JW, Roberson EDO, Anbunathan H, Onken MD, Worley LA, Bowcock AM. Recurrent mutations at codon 625 of the splicing factor SF3B1 in uveal melanoma. Nat Genet. 2013;45:133–5.

40. Sevostyanova IA, Kulikova KV, Kuravsky ML, Schmalhausen EV, Muronetz VI. Sperm-specific glyceraldehyde-3-phosphate dehydrogenase is expressed in melanoma cells. Biochem Biophys Res Commun. 2012;427:649–53.

41. Dobin A, Davis CA, Schlesinger F, Drenkow J, Zaleski C, Jha S, et al. STAR: ultrafast universal RNA-seq aligner. Bioinformatics. 2013;29:15–21.

42. Cerami E, Gao J, Dogrusoz U, Gross BE, Sumer SO, Aksoy BA, et al. The cBio cancer genomics portal: an open platform for exploring multidimensional cancer genomics data. Cancer Discov. 2012;2:401–4.

43. Widmann J, Stombaugh J, McDonald D, Chocholousova J, Gardner P, Iyer MK, et al. RNASTAR: An RNA STructural Alignment Repository that provides insight into the evolution of natural and artificial RNAs. RNA. 2012;18:1319–27.

44. Nelder JA, Mead R. A Simplex Method for Function Minimization. Comput J. 1965;7:308–13.

