## Supplemental Figures for "Bisbee: A proteomics validated analysis package for detecting differential splicing, identifying splice outliers, and predicting splice event protein effects"

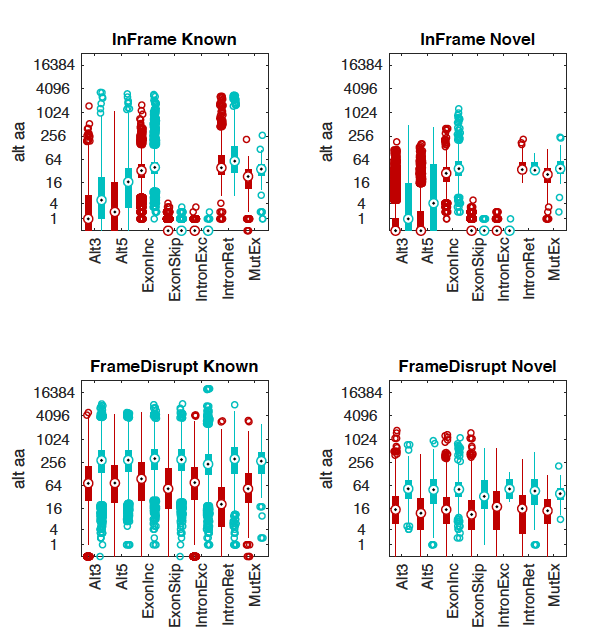

##### Supplemental Figure 1. Length of altered amino acid sequence generated by splice events

*Boxplots illustrate distribution of the length of amino acid sequence differing between the two isoforms for splice events on a log scale. Red bars show the distribution for all of the predicted events and blue bars show the distribution only for events with confirmed protein expression for each event type. As expected, events generating longer altered regions are more likely to be detected in mass spectrometry data. Events involving wild-type isoforms tend to generate longer affected regions than those generating novel isoforms.*

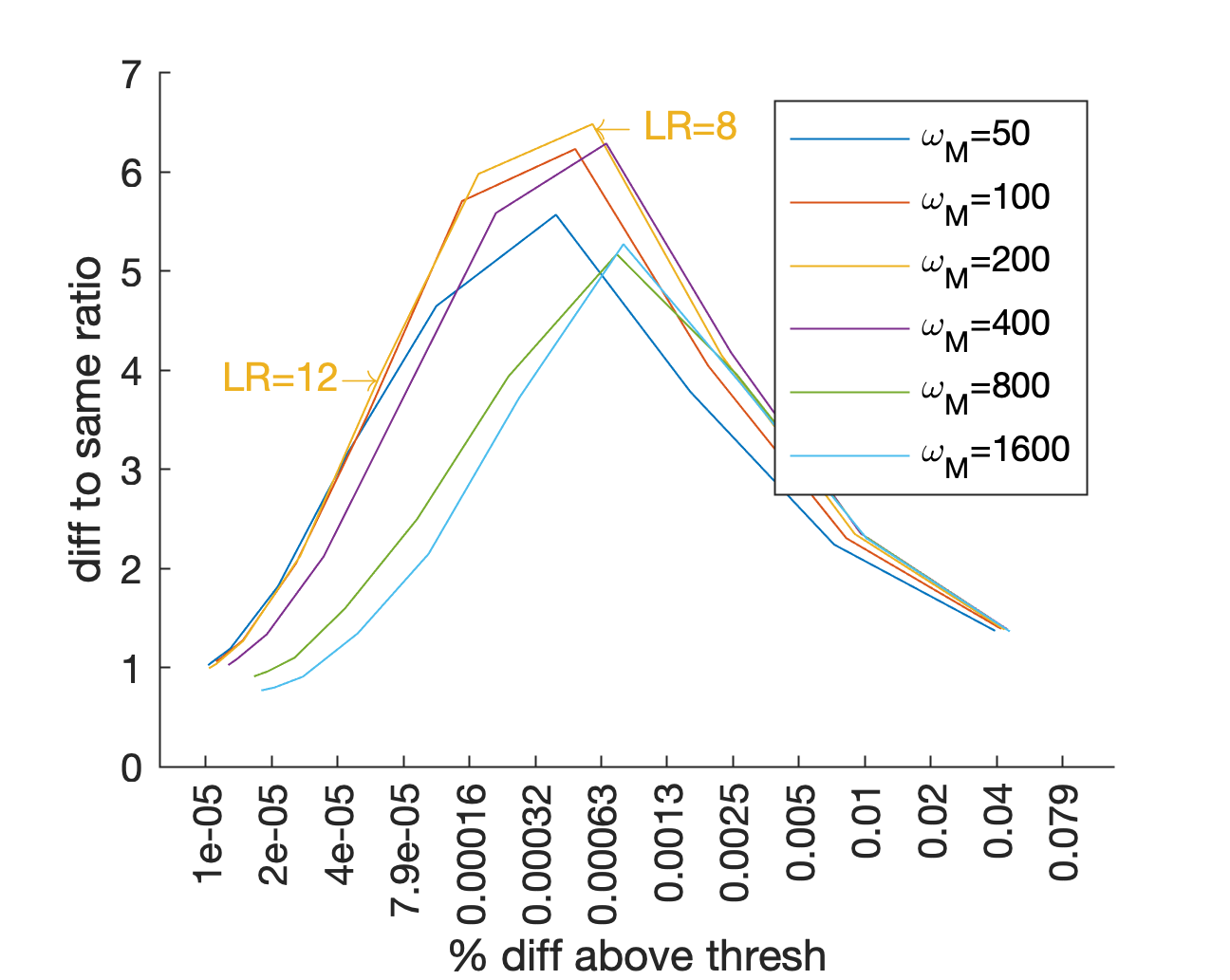

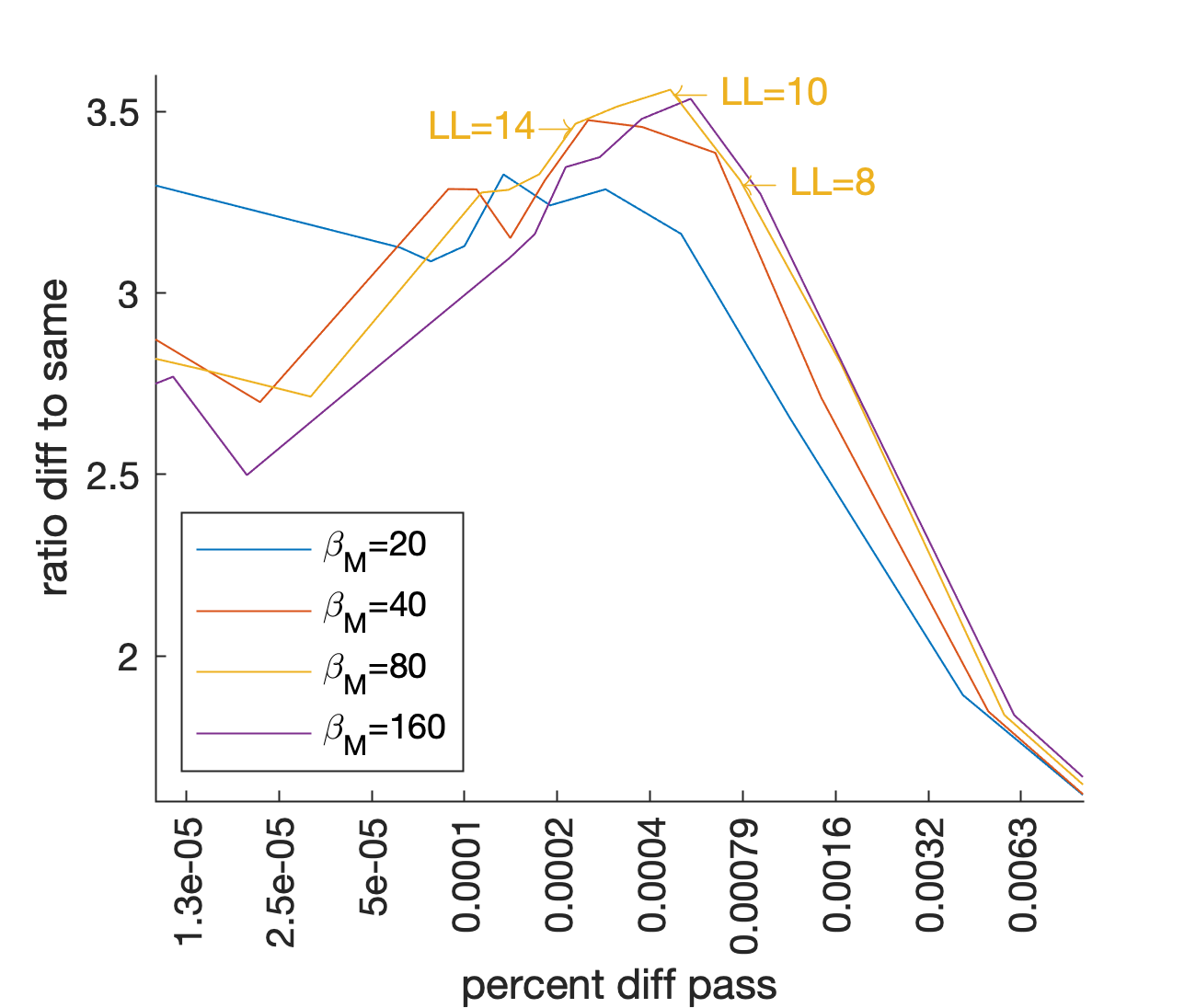

##### Supplemental Figure 2. Parameter optimization.

*a) The differential beta binomial test was run on random sets of samples from the same tissue or different tissues selected from GTEx with different values of the* $\omega_{M}$*parameter. The ratio of percentage of events that are higher than a given LR threshold in the different tissue comparison compared to the same tissue comparison is plotted against the percentage of events in the different comparison that pass the threshold. b) The outlier model was trained on 12 different tissues with 80 samples in each set using different values of the maxW parameter. Outlier scores were found for each of the 12 models for a different set of test samples selected from the same tissues. The ratio of the percent of data points passing an outlier score threshold between models fit on the same tissue vs different tissues is compared to the percentage of data points from fitting unmatched tissue models that pass the threshold.*
